# The Noise-Resilient Brain: Resting-State Oscillatory Activity Predicts Words-In-Noise Recognition

**DOI:** 10.1101/705053

**Authors:** Thomas Houweling, Robert Becker, Alexis Hervais-Adelman

**Author notes:** Correspondence to., Neurolinguistics, Department of Psychology, University of Zürich, Binzmühlestrasse 14, box 5, CH-8050, Zürich, Switzerland.

## Abstract

The role of neuronal oscillations in the processing of speech has recently come to prominence. Since resting-state (RS) brain activity has been shown to predict both task-related brain activation and behavioural performance, we set out to establish whether inter-individual differences in spectrally-resolved RS-MEG power are associated with variations in words-in-noise recognition in a sample of 88 participants made available by the Human Connectome Project. Positive associations with resilience to noise were observed with power in the range 21 and 29Hz in a number of areas along the left temporal gyrus and temporo-parietal association areas peaking in left posterior superior temporal gyrus (pSTG). Significant associations were also found in the right posterior superior temporal gyrus in the frequency range 30 to 40Hz. We propose that individual differences in words-in-noise performance are related to baseline excitability levels of the neural substrates of phonological processing.

**Highlights:**

- Power of resting MEG activity predicts Words-In-Noise recognition performance
- Significant associations in higher beta and lower gamma frequency band
- Strongest in left-lateralised perisylvian cluster peaking in posterior STG
- Effects are spectrally and spatially consistent with phoneme-level processing

## 1. Introduction

Over the last decade, the scope of neuroscientific inquiry has considerably broadened, as the traditional approach to studying the brain through its task-elicited activation, is increasingly assisted and complemented by the interrogation of its task-independent properties (Biswal et al., 2009). Among the factors contributing to the rise in interest in spontaneous brain activity has been the realisation that resting-state (RS) brain activity has the potential to index a number of intrinsic features of the brain. Further, measures of RS brain activity have been shown to be predictive not only of task-related brain activation (Cole, Ito, Bassett, & Schultz, 2016; Mennes et al., 2010, 2011; Tavor et al., 2016), but also of behavioural outcomes (Finn et al., 2016; Fox, Snyder, Vincent, & Raichle, 2007; Fox, Snyder, Zacks, & Raichle, 2006; Mennes et al., 2011). Contrary to traditional hypothesis-driven task-based approaches that aim at characterising how – where and when – the brain processes the given stimulus property, as indexed by brain activation, hypothesis- and task-free approaches aim to characterise inherent features of the brain that shape how information from the outside world is processed. To this extent, the brain’s RS activity may be conceived of as a structuring context in which evoked activity occurs.

In this study we used magnetoencephalography (MEG) to characterise the brain’s RS activity in terms of spectrally-resolved power. MEG records magnetic field perturbations produced by neuronal electrical activity at high temporal resolution, enabling the decomposition of the broadband signal into spectrally-resolved components. It has been shown that different frequency ranges are functionally distinct (Buzsáki & Draguhn, 2004). Spectrally-specific modulations in amplitude/power of the electromagnetic field associated with neuronal activity are thought to reflect the degree of synchronisation among neuronal populations. Here, we tested whether individual differences in spectrally-resolved MEG power relate to inter-individual variability in a speech perception task in a sample of 88 participants made available as part of the Human Connectome Project (HCP; Van Essen et al., 2013). Specifically, we tested whether spectrally-resolved RS MEG power is predictive of the ability to perceive speech in noise, as measured by a Words-In-Noise (WIN) test. WIN assesses individuals’ ability to recognise isolated monosyllabic words embedded in background noise. The ability to comprehend speech in noise, and acoustically-degraded speech more generally, is known to be highly variable across individuals (Mattys, Davis, Bradlow, & Scott, 2012), and several hypotheses have been advanced regarding cognitive determinants of speech in noise and degraded speech comprehension (Rönnberg et al., 2013; Zekveld, Rudner, Johnsrude, & Rönnberg, 2013). Here, however, we aimed to examine the relationship between RS activity and WIN performance, while controlling for a battery of other, general cognitive and demographic factors. In this way, we hope to make progress in elucidating the neural bases of speech-in-noise comprehension that are determined by intrinsic brain properties, dissociated from cognitive factors.

It is thought that a first step in the analysis of sensory signals is their sampling and quantisation. Cortical oscillations, which are rhythmic fluctuations in neuronal synchrony at specific time scales, have been proposed as a key mechanism responsible for the processing of the incoming continuous speech stream. According to the influential asymmetric sampling in time hypothesis (AST; Giraud et al., 2007; Giraud & Poeppel, 2012; Poeppel, 2003), a “*principled relation*” (i.e., a quasi-isomorphism) exists between the time scales at which meaningful acoustic events in speech occur and those at which the speech perception system operates, by means of cortical oscillations. This hypothesis holds that speech processing is temporally-asymmetric in non-primary auditory areas, with left-hemisphere (LH) mechanisms (i.e., low-γ oscillations in the ∼20-50Hz frequency range) preferentially extracting information over shorter (20-50ms) temporal windows and right-hemisphere (RH) mechanisms (i.e., θ oscillations in the ∼3-7Hz frequency range) over longer (∼150-300ms) windows (Poeppel, 2001, 2003). The slower time scale corresponds to the duration of slow spectrotemporal fluctuations associated with the alternation of syllables and the latter corresponds to a processing or sampling rate suitable for capturing the rapid acoustic features relevant for phonemic identification, such as formant transition and voice onset time (Delattre, Liberman, & Cooper, 1955; Kewley-Port, 1982; Liberman, Cooper, Shankweiler, & Studdert-Kennedy, 1967; Lisker & Abramson, 1964; Rosen, 1992).

The AST suggests that the evident functional asymmetry of non-primary auditory cortices is related to differences in the distribution of the centre frequency at which neuronal ensembles spontaneously synchronise. These are relatively more skewed towards synchronising at a θ rate in the RH and skewed towards synchronising at a low-γ pace in the LH. There is indeed some evidence for spectral peaks in θ and low-γ frequency ranges in resting electrophysiological activity within primary auditory areas, as observed, for instance, by Lakatos and collaborators (2005) by means of intracranial recordings in macaques. These spontaneous, ongoing oscillations were also shown to predict the activity evoked by noise bursts. Further, consistent with the slight asymmetry in spontaneous activity implied by the AST, it has been shown by means of simultaneous EEG and fMRI recordings in humans that spontaneous activity in the low-γ range at rest correlates best with LH auditory cortical synaptic activity (as indexed by the BOLD signal), whilst RS EEG power within the θ range correlates best with synaptic activity in the RH (Giraud et al., 2007). Finally, Lehongre and collaborators (2011) have shown asymmetries in peak oscillatory responses to rhythmic auditory stimulation in planum temporale and posterior superior temporal sulcus: whilst stronger responses where elicited in the RH by periodicities above 55Hz, the strongest responses in the LH were elicited by periodic auditory stimulation in the 25-35Hz range.

We propose that individual differences in WIN performance may be related to baseline excitability of the speech perception system. While there is evidence for a number of oscillatory regimes to be involved in the processing of language – with θ and low-γ being of paramount interest – evidence that the amplitude of spectrally-resolved spontaneous activity is able to predict a rather high-level speech perception task as WIN recognition has not been reported thus far. To date, the wide majority of studies on task-free brain activity have focused on examining its functional connectivity – i.e., the temporal correlation between spatially remote brain areas which is thought to represent communication – by means of functional magnetic resonance imaging (fMRI). By measuring coherent spontaneous low-frequency (i.e., <0.1Hz) fluctuations of the blood oxygenation level dependent signal, studies making use of fMRI have demonstrated the existence of a number of large-scale neural networks, which highly resemble the activation and deactivation maps associated with performing perceptual, motor or cognitive tasks (see Van Den Heuvel & Pol, 2011 for a review). Here, in contrast, we use a relatively simple representation of the brain’s RS characteristics in a bid to predict a specific behavioural ability. Importantly, rather than measuring information flow over large and distributed networks, as would be the case in connectivity analyses, we aimed at indexing local brain activity and how it relates to individual behavioural differences.

## 2. Material and methods

### 2.1. Participants and procedures

Resting-state MEG data from 88 participants were analysed. Mean age was 28 years and 7 months (SD = 4 years, range = 22-35 years). Forty-one participants were female. All of the subjects had normal hearing. The full HCP protocol of participants undergoing MEG is typically completed in a three-day visit (see the Reference Manual for the 1200 Subjects Data Release by the WU-Minn Consortium Human Connectome Project, 2017 for details: https://www.humanconnectome.org/study/hcp-young-adult/document/1200-subjects-data-release). RS MEG data acquisitions always preceded WIN test by at least one full day.

### 2.2. Speech material

The Words-In-Noise test (WIN; Wilson, 2003; Wilson et al., 2007) was used to test participants’ ability to recognise single words presented amid varying levels of background noise. The WIN establishes individual signal-to-noise ratio (SNR) thresholds for correct word recognition and, by so doing, measures how much difficulty a person might have listening to speech in a noisy environment. Specifically, participants are presented with isolated monosyllabic words embedded in varying levels of multitalker babble. They are asked to listen to and then repeat the words. The task becomes increasingly difficult as the background noise gets louder, thus reducing the SNR. Multitalker babble involves several (i.e., six; three male and three female) speakers talking simultaneously about various topics with all of the conservations being unintelligible (Sperry, Wiley, & Chial, 1997). Multitalker babble is the most common environmental noise encountered by listeners in everyday life and is, therefore, highly ecologically valid (Plomp, 1978). Furthermore, it is more sensitive to mild hearing loss as compared to speech-spectrum noise (Findlay, 1976).

Thirty-five words are presented in blocks of five items whilst SNR decreases monotonically through seven levels (26, 22, 18, 14, 10, 6, 2 and -2 dB SNR). The test ends when all the seven lists of items have been completed or when all the five items of a given SNR are not correctly recognised. The examiner assigns one point for each item correctly recognised and no points to incorrect responses. The best raw score a participant can obtain is 35 (all correct) and the worst is zero (none correct). Raw scores are converted back to dB SNR thresholds via the formula: *SNR threshold (dB) = 26 dB -.8 * raw score.* SNR thresholds can therefore take values from 26 dB (none correct) to -2 dB (all correct). In the sample we analysed SNR thresholds ranged from 7.6 to 2 dB (mean=4.7, SD=1.3). Of note, participants composing the current sample have been tested through either one of two versions of the NIH Toolbox WIN test (V1 and V2). Participants tested through V1 had stimuli delivered via speakers, whilst the majority of V2 participants had stimuli delivered monaurally through headphones, and the better score (lowest SNR) out of the two ears was used as participant score. V1 and V2 scores were normed to be directly comparable across the whole 1200-subjects release. We verified this is the case in our limited subset of 88 participants by testing for mean differences across WIN score in the two groups using the independent 2-group Mann-Whitney U Test (two-way). The test revealed that participants tested through WIN V1 (median = 5.2 dB SNR) did not perform differently from participants tested with VS2 (median = 4.4 dB SNR), W=846, *p*=.939. Note, however, that some of the participants labelled as ‘V2’ could have actually been tested with WIN V1.

#### 2.2.1. Deconfounding procedure

We aimed to establish whether resting MEG power is specifically associated with the ability to recognise words in noise. However, several unspecific factors and variables including demographics (e.g., age) and general cognitive abilities (e.g., working memory, processing speed), may contribute to (i.e., confound) each participants’ SNR threshold (Frisina & Frisina, 1997; Humes et al., 1994). For this reason, we exploited the wide range of participant measures and information collected as part of the HCP to control for potential contributions of these factors. A detailed description of the measures collected as part of the assessment is available in Van Essen and collaborators (2013). In short, these measures include demographic information, neurological/psychiatric diagnoses, description of subjects’ habits (e.g., sleep, drug consumption, etc.), and a wide range of perceptual, cognitive and emotional tests and questionnaires. Out of the full set of subject measures (reported in the “open access” and “restricted” subject information spreadsheets available here: http://humanconnectome.org/data), we selected a subset of 211 variables. The full list of these variables can be found in the Appendix. We then performed a second selection of those measures meeting the criteria outlined in Table 1 (adapted from Smith et al., 2015). This procedure resulted in the selection of a final set of 178 confounds.

**Table 1.** Criteria for the selection of subject measures entering the PCA (adapted from Smith et al., 2015).

| Criterion | Formal requirement |
| --- | --- |
| Subject measure data need to have enough variability | 1. Standard deviation ($SD$) > 0 |
|  | 2. At least 49 valid subject scores |
|  | 3. The number of values in the largest group of unique values must not exceed 80% of the total number of valid values |
| Subject measure data need to have excessively high leverage points | 4. If $x_s$ is a subject measure value for subject $s$, and $y_s = (x_s - \text{median}(x_s))^2$, the subject measure must not contain any $x_s$ for which $\max(y_s) > 100 \times \text{mean}(y_s)$. |

Of these measures, 20 were found to be significantly correlated with WIN SNR after FDR correction for multiple comparisons (Fig. 1). Specifically, seven measures of tobacco consumption and dependence were negatively associated with resilience to noise (positively associated with WIN SNR; *r*=.243 – .352, *p*_*(FDR)*_=.047 – .013). This is in line with a large population-based study reporting that, compared to non-smokers, smokers are more likely to experience difficulties in speech-in-noise understanding in a dose-dependent fashion (Dawes et al., 2014). In addition, also three indices of social withdrawal and perceived social rejection negatively predicted resilience to noise (*r*=.244 – .262, *p*_*(FDR)*_=.048 – .042), in line with the reported association between hearing difficulties and social isolation (Ciorba, Bianchini, Pelucchi, & Pastore, 2012; Weinstein & Ventry, 1982). Further, ten measures positively predicted resilience to noise (negatively correlated with WIN SNR), namely both age-adjusted and -unadjusted scores on NIH flanker task (*r=-.*354, *p*_*(FDR)*_=.027, and *r=-.*348, *p*_*(FDR)*_=.016, respectively), both age-adjusted and -unadjusted scores on NIH card sorting test (*r=-.*330, *p*_*(FDR)*_=.013, and *r=-.*317, *p*_*(FDR)*_=.016, respectively), both age-unadjusted and -adjusted scores on NIH list sorting test (*r=-.*306, *p*_*(FDR)*_=.017, and *r=-.*303, *p*_*(FDR)*_=.017, respectively), both age-adjusted and unadjusted score on NIH dexterity test (*r=-.*255, *p*_*(FDR)*_=.047, and *r=-.*242, *p*_*(FDR)*_=.048, respectively), education level (*r=*-.279, *p*_*(FDR)*_=.009), and in-fMRI-scanner accuracy on the language (‘Story’) task (*r=*-.234, *p*_*(FDR)*_=.026). It is interesting to note that (verbal) working memory and executive functions have been highlighted as cognitive contributors to degraded speech comprehension (Rönnberg et al., 2013) and that response inhibition seems to be particularly required to perceive speech-in-noise when noise is verbal as compared to non-verbal (Rouleau & Belleville, 1996), as is the case in the context of the WIN test (multitalker babble). With respect to education, we speculate that it may help compensating for a lower perceptual quality of the auditory object in individuals with sub-optimal (although still normal) hearing-in-noise. Finally, with regards to the in-scanner performance on the language task, it is clear that higher resilience to scanner noise may allow for a clearer perception and thus comprehension of the story to be heard. We nonetheless regressed out every available variable, including those that were not significantly individually associated with the measure of interest because the subject measures taken together are sources of individual differences that we wished to remove, as we aimed at establishing relationships that are specific to WIN (See Fig. S1-S2 for an overview on how scores changed as a result of the deconfounding procedure). In order to avoid overfitting in the regression, principal component analysis (PCA) was used to reduce the dimensionality of the set of confounds. Before entering the PCA, variables were gaussianised by means of rank-based inverse normal transformation and standardised (to mean=0 and SD=1), by subtracting the sample mean from each individual score and dividing the demeaned score by the standard deviation. The statistical rationale for normalising confounds prior to PCA is to stabilize the variance (reduce heteroscedasticity) and to facilitate its quantification and comparison across the range of measures. Twenty-five principal components, accounting for 80% of the total variance of the set of 132 measures, were regressed out from the original WIN score following the steps described by Smith and collaborators (2015).

**Figure 1.**
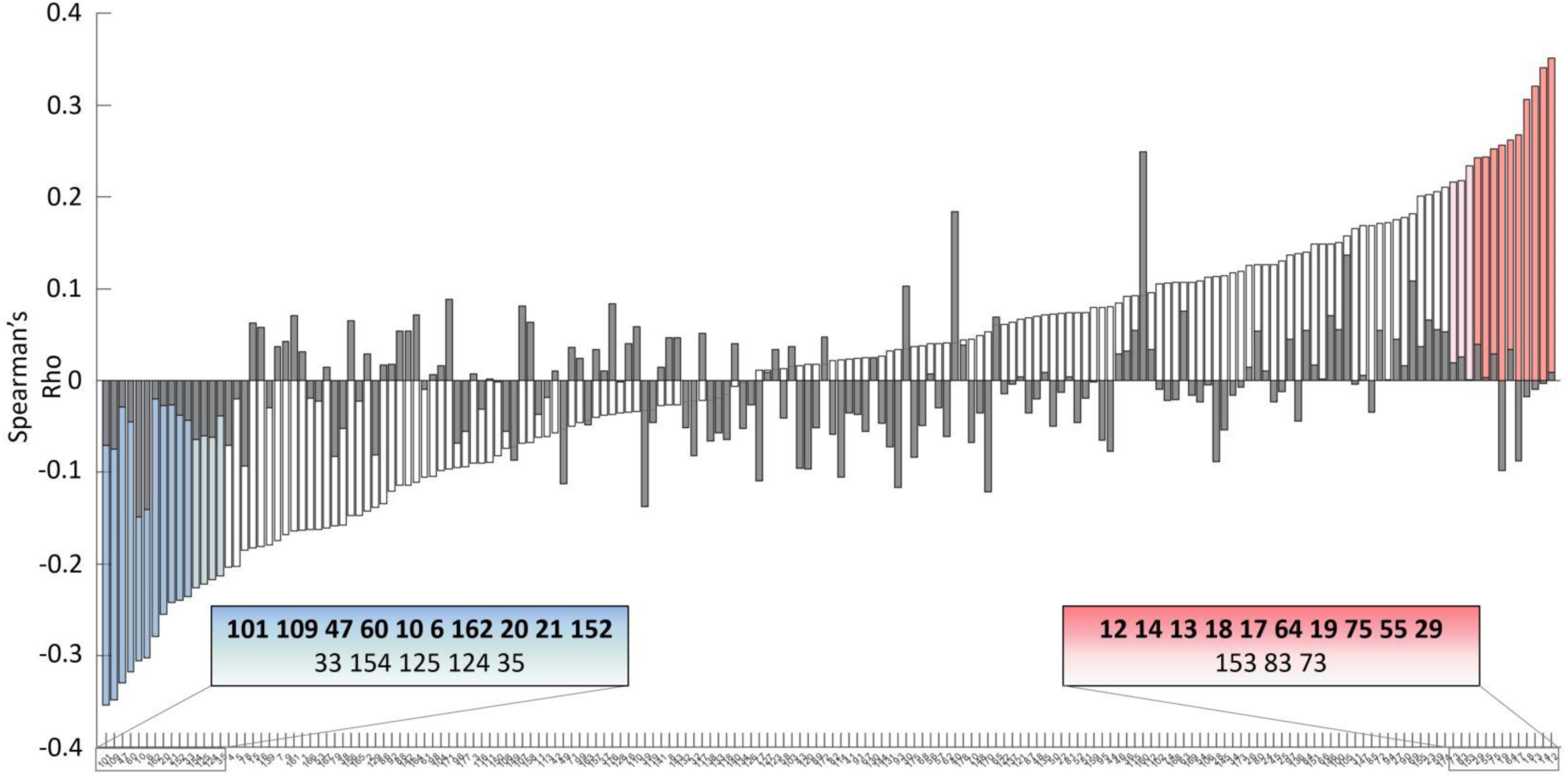
Effect of deconfounding WIN scores on the association with confounds. X-axis represents confounds (one per bar, sorted ascendingly by correlation coefficient; numbers represent the confound label ‘ID_out’ in Appendix) and Y-axis represents the correlation coefficient (Spearman’s Rho) between WIN SNR and each confound. Color-codes: white bars = non-significant correlations between original (non-deconfounded) WIN SNR scores and confound; light-blue bars = significant (p<.05, uncorrected) negative correlations between original WIN SNR scores and confound; dark-blue bars = significant (p<.05, FDR-corrected) negative correlations between original WIN SNR scores and confound; pink bars = significant (p<.05, uncorrected) positive correlations between original WIN SNR scores and confound; red bars = significant (p<.05, FDR-corrected) positive correlations between original WIN SNR scores and confound; overlayed grey bars = (non-significant) correlations between deconfounded WIN SNR and confound. Numbers in bold within boxes represent the confound label of significant FDR-corrected WIN-confound correlations, whilst the others represent the confound label of significant non-corrected WIN-confound correlations.

### 2.3. RS-MEG data

RS-MEG data was acquired in three runs of approximately six minutes per subject during which participants are asked to remain supine with eyes open. Details on acquisition protocols and hardware specifications can be found in in the 1200 Subject Data Release Manual linked above. Briefly, subjects were scanned on a whole head MAGNES 3600 (4D Neuroimaging, San Diego, CA) system housed in a quiet, darkened and magnetically shielded room. The MEG system includes 248 magnetometer channels together with 23 reference channels. Data sampled at 508.63 Hz were downloaded from https://db.humanconnectome.org/. We applied a fifth-order Butterworth low-pass filter at 48Hz (by means of FieldTrip’s ‘ft_preproc_lowpassfilter’ implementation) and down-sampled the data to 200 Hz by means of Matlab’s ‘resample’ routine with default input parameters. Raw MEG data underwent the standard HCP MEG data pre-processing pipeline based on independent component analysis (ICA) aimed at identifying and removing artefacts, bad channels and bad segments (2s each). Additionally, single shell volume conduction models (one per participant) were defined in the MEG-system based head coordinate system from segmentation of anatomical MRI T1 images. We computed the inverse solution using Linearly Constrained Minimum Variance (LCMV) Beamformer filters. These allow source models to be defined on a regular 3D grid in normalized MNI-space with a resolution of 8mm, aligning the subjects in source space. Welch’s method was used to obtain the power spectral density (PSD) of each of the source-reconstructed and standardised (over time to mean=0 and SD=1) time series session data (three per participant, mean duration = 281s). The two-sided PSD was computed on eight equally-long time series windows of approximately 62.5s each with 50% overlap from 1 to 40 Hz (in 1-Hz bins) at each source point (3559 voxels of size 8*8*8mm each). PSDs of the three sessions were then averaged within participants for each voxel.

#### 2.3.1. Definition of frequency bands

In order to accommodate the inherent similarity of the power distribution in neighbouring frequencies, *k*-means clustering (1000 replicates, using correlation distances) was used to cluster the 40 frequency bins into 6 wider bins based on the similarity of the spatial distribution of voxelwise power. Evaluation of the optimal clustering solution was based on a two-step procedure. First, whole-brain voxelwise power cross-correlation matrix was inspected visually in order to determine a range of *k*s to test. Once the range of clusters to evaluate was established (4 to 6), the optimal solution was determined using two measures of internal clustering validation measures, namely Dunn’s Index (DI; Dunn, 1973), and Silhouette Index (SI; Rousseeuw, 1987). Both indices indicated that *k*=6 represented the best clustering solution. The final frequency bins – mainly corresponding to conventional frequency bands – were as follows: 1-4Hz (‘d’), 5-7Hz (‘θ’), 8-15 (‘α’), 16-20 (‘low-β’), 21-29Hz (‘high-β’), and 30-40Hz (‘low-γ’).

### 2.4. Statistical Analyses

Within-subject means of voxelwise power in each frequency band were correlated with the standardised deconfounded WIN score. We evaluated the statistical significance of the clusters by means of permutation testing. Because the sample analysed here is composed by subgroups of monozygotic and dizygotic siblings in addition to non-related participants, we took into account the possible effect of family structure on the null distribution of correlations generated by permutation by randomly shuffling WIN scores (10000 times) across participant labels in a manner constrained by family structure. Multi-level block permutation (Winkler, Webster, Vidaurre, Nichols, & Smith, 2015) was performed using FSL’s PALM utility (https://fsl.fmrib.ox.ac.uk/fsl/fslwiki/PALM), and the exchangeability block structure specified in the HCP documentation (https://fsl.fmrib.ox.ac.uk/fsl/fslwiki/PALM/ExchangeabilityBlocks). For each frequency band (n=6) and voxel (n=3559), a *t*-value was calculated for both the actual and the data-driven null distribution; *t*-values were then compared to produce *p*-values (separately for the positive and negative contrasts). Below we report results that are significant at a voxelwise level of *p*<.05, corrected for family-wise error rate (FWER) of the multiple comparisons over the 3559 solution points.

## 3. Results

RS activity was found to be predictive of WIN performance in two adjacent frequency ranges (Table 2, Figure 2). In both the high-β (21-29Hz) and low-γ (30-40Hz) bands, we observed significant negative correlations between power and WIN SNR, indicative of a positive association between amplitude of the oscillatory activity and the ability to be resilient to noise-induced breakdown of intelligibility. In the high-β frequency band, significant associations clustered around four peaks. The largest and most significant peaks are found in the posterior section of the superior and middle temporal gyri (STG and MTG, respectively) and in a more anterior section of the MTG. We further observed small clusters of correlation in associative areas within the lateral aspect of the middle occipital gyrus and inferior parietal lobule (two and one voxels, respectively). In the low-γ band, we notice a cluster of correlation in the RH in the posterior part of the STG.

**Table 2.** Summary of loci of significant correlations between RS brain activity and WIN recognition. Anatomical labels were obtained using in-house routines to assign identity according to the nearest labelled coordinate in the AAL template (Tzourio-Mazoyer et al., 2002), Brodmann area labels were extracted from the BA template provided with MRIcron (https://www.nitrc.org/projects/mricron). Morethan one peak per cluster is reported if peaks are at least 24mm apart.

|  | Cluster ID | N<br>vox | AAL Label | BA | x,y,z | t | p |
| --- | --- | --- | --- | --- | --- | --- | --- |
| **High-β** | 1 | 24 | Left superior temporal gyrus | BA42 | -62,-42,16 | 2.602 | 0.002 |
|  |  |  | Left middle temporal gyrus | BA37 | -62,-66,0 | 2.06 | 0.009 |
|  | 2 | 15 | Left middle temporal gyrus | BA21 | -62,-18,-16 | 2.102 | 0.008 |
|  | 3 | 2 | Left middle occipital gyrus | BA18 | -46,-90,-8 | 1.556 | 0.028 |
|  | 4 | 1 | Left inferior parietal gyrus | BA40 | -46,-58,56 | 1.383 | 0.041 |
| **Low-γ** | 1 | 3 | Right superior temporal gyrus | BA48 | 66,-42,24 | 1.602 | 0.025 |

**Figure 2.**
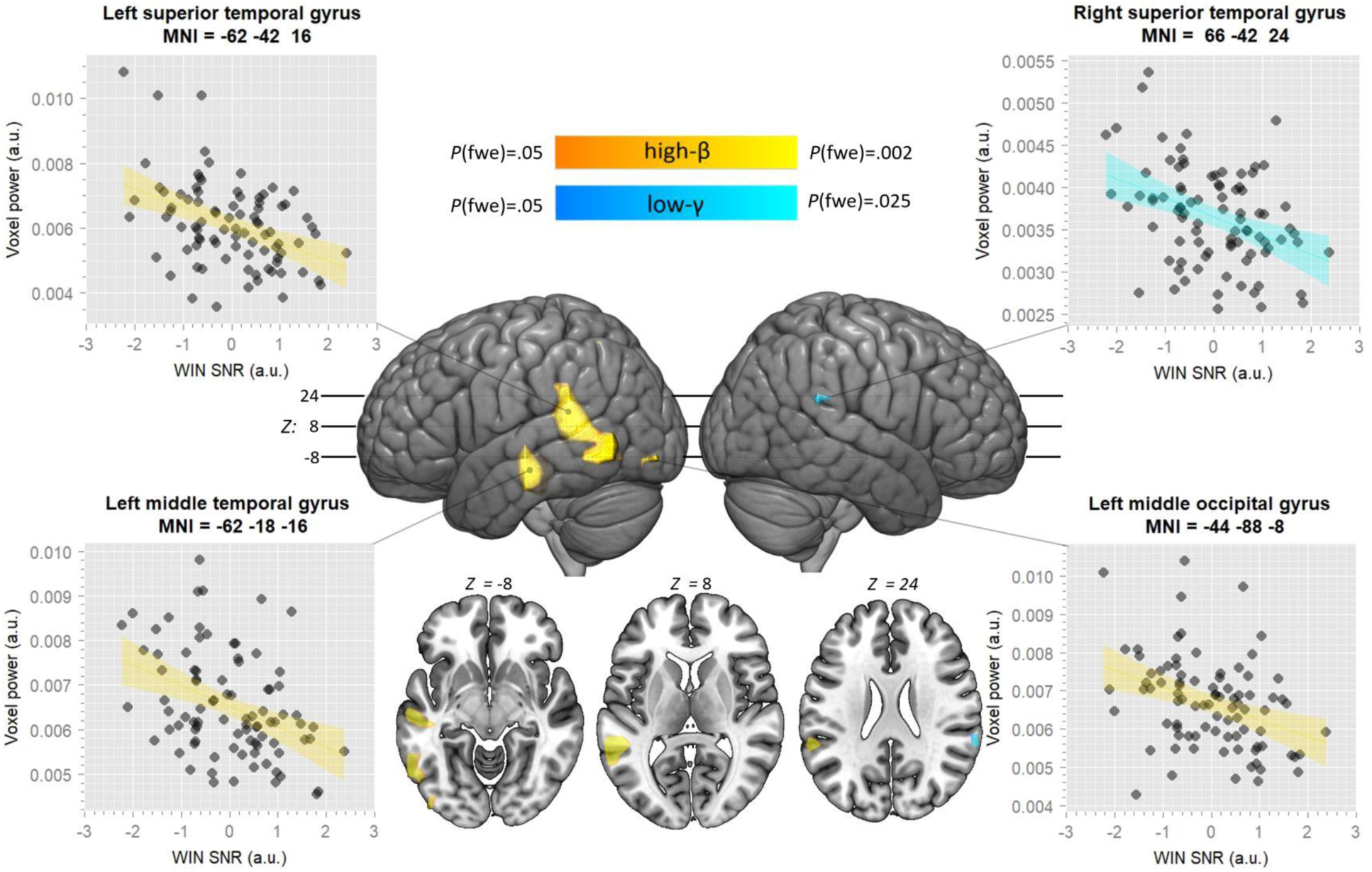
Surface rendering and axial slices of significant correlations between WIN SNR and voxel power. Significant correlations in the high-β band are represented in warm colours, whilst significant associations in the low-γ band are represented in cold colours. The scatterplots represent the association at each of four representative voxels by means of a regression line with standard error bounds.

## 4. Discussion

This analysis has revealed that topographically- and spectrally-resolved RS MEG power – an index of spontaneous neuronal synchronisation at various time scales – can predict behavioural performance. This analytical approach offers an alternative and complementary perspective on mechanisms of brain functioning as compared to activation studies. Indeed, whilst in the latter type of studies, individual differences in brain responses to stimulation are generally treated as random noise and often ascribed to confounding ‘volatile’ factors, such as participants’ momentary alertness level, here we specifically tested the functional relevance of such differences. Importantly, the observed effects cannot be accounted for by a spurious association with volatile confounds, since the electrophysiological and behavioural assessments took place in different contexts, separated by a day or more. Furthermore, we removed the potential confounding effects of a substantial array of cognitive and demographic factors, ensuring that the results we report are specific to the relevant behavioural domain, namely speech processing. In attempting to remove confounding factors from the data we also revealed how WIN recognition performance is associated with a number of indices (Fig. 1). Although a comprehensive discussion of such associations is beyond the scope of the present report, one that is arguably of special interest is that with verbal working memory (vWM). vWM has long been implicated in speech-in-noise recognition (Rönnberg et al., 2013), though a recent meta-analysis has concluded that this relation only holds true in the context of aging and/or hearing impairment (Füllgrabe & Rosen, 2016). Here, intriguingly, we observed this association in a sample of young normally-hearing participants.

The results do not appear to shadow the left-right asymmetries of the AST, whereby LH auditory regions are posited to express relatively more 20-50Hz activity than RH auditory areas that express greater lower-frequency power in response to speech input. Although potentially surprising at first glance, the AST describes responses to speech input, without making any specific predictions about the system at rest, beyond the implication that it is tuned to engage with acoustic information at the proposed timescales. The spatial distribution of the present results seems to indicate that individual differences in WIN performance relate to baseline excitability levels of a number of non-primary areas in the LH known to support speech recognition that are involved in the processing (left posterior STG/STS) of phonological information and its integration with lexical (left posterior/middle MTG) information, as well as with somatosensory input from other modalities (left IPL and visual association areas; Friederici, 2012; Hickok & Poeppel, 2007). Thus, we suggest that increases in spectrally- and topographically-specific MEG power at rest, by determining a suitable context in which relevant acoustic features (i.e., cues to phonemic identity) are processed, are associated with better WIN recognition abilities.

This proposal is consistent with a number of lines of evidence. The locus of the LH high-β peak is consistent with the region in which categorical phoneme perception has been demonstrated in fMRI (e.g., Specht, Baumgartner, Stadler, Hugdahl, & Pollmann, 2014; see Turkeltaub & Branch Coslett, 2010 for a review). Evidence from intracranial recordings also supports this. For example, Chang and colleagues (2010) carried out intracranial recordings of STG responses to synthesised phonemic continua and found neural populations in pSTG responding to phonemic cues important for phoneme recognition. In addition, they observed populations of neurons responding selectively and categorically to single phonemes, providing a strong physiological correspondence to the behavioural categorical perception phenomenon. A further study (Mesgarani, Cheung, Johnson, & Chang, 2014) has shown that neuronal populations discretely responding to phoneme categories are hierarchically clustered in a way that is strikingly similar to the traditional clustering of phonemes performed by phoneticians and linguists. Mesgarani and colleagues (2014) conclude that selectivity is determined primarily by acoustic cues for manner and secondarily by place of articulation (and, in general, by articulatory properties). Further, we observe areas of significant WIN recognition – RS power association more anteriorly along the left STG and MTG. These areas have been implicated by a number of studies in the representing phonemes with higher specificity and invariance (Ashtari et al., 2004; DeWitt & Rauschecker, 2012; Liebenthal, Binder, Spitzer, Possing, & Medler, 2005).

Brain-behaviour relationships observed in this study were confined to two neighbouring frequency bands (21-29 and 30-40Hz). Different lines of inquiry indicate that neural activity within this range of the spectrum is important for the parsing and processing of acoustical information on short time scales relevant for phoneme recognition, i.e., phonetic features. Among the work supporting this hypothesis, relatively strong evidence comes from a growing body of research on exogenous modulation of neural excitability. Recent studies using transcranial alternating current stimulation (tACS) have begun to provide causal evidence for the role of oscillations in this frequency range for phoneme perception. The oscillatory current produced by tACS is assumed to instantaneously synchronize (i.e., to entrain) the firing in the targeted neuronal populations, enhancing oscillatory power at the stimulation frequency (Helfrich et al., 2014). A recent study using tACS reported the induction of phoneme recognition deficits in healthy participants as a result of applying a 40Hz current to auditory areas (Rufener, Zaehle, Oechslin, & Meyer, 2016). In a separate study, Rufener, Zaehle and colleagues (2016) found that applying 40Hz stimulation to the bilateral auditory cortex reduced the rate of perceptual learning during a phoneme categorisation task, while applying 6Hz tACS stimulation did not affect perceptual learning. Importantly, whilst this result was replicated in a comparable sample of young participants (20-35 years), it was shown that perceptual learning of older participants (60-75 years) was enhanced, rather than hindered, by 40Hz stimulation and the presumed tACS-induced enhancement of 40Hz activity, with 6Hz stimulation again having no significant effect (Rufener, Oechslin, Zaehle, & Meyer, 2016). Intriguingly, Rufener and colleagues suggest that the effect of perturbing low-γ activity may depend on (i.e., interact with) its ‘pre-existing’ (i.e., RS) level.

The idea that there may be such a thing as an optimal resting-state level for phoneme processing is further supported by experiments on developmental dyslexia (DD). DD is a common disability defined as sub-normal acquisition of reading skills and phonological abilities within the context of otherwise normal hearing and cognitive abilities. Rufener and colleagues (Rufener, Krauel, Meyer, Heinze, & Zaehle, 2019) showed that 40Hz tACS stimulation over the bilateral auditory cortex improved phoneme-categorisation accuracy in a sample of patients suffering from DD. Thus, by altering the putatively optimal balance of synchronous endogenous activity in healthy individuals unaffected by phonological deficits, sampling of rapid acoustic events is altered and performance is worsened. In contrast, when the equilibrium of oscillatory activity is sub-optimal, as might be the case in the context of healthy aging (Voytek et al., 2015) and DD (Hancock, Pugh, & Hoeft, 2017), abnormal, or sub-optimal, temporal sampling is remediated by the external stimulation, and performance improves. Taken together, evidence from these tACS studies strongly supports a causal involvement of low-γ activity in the rapid auditory processing hypothesised to be essential for phoneme extraction and identification.

A large body of evidence suggests that the phonological deficits that characterise DD may arise from abnormal sampling of the speech stream into the short time windows relevant for the processing of phonetic cues (Chobert, François, Habib, & Besson, 2012; Gaab, Gabrieli, Deutsch, Tallal, & Temple, 2007; Raschle, Stering, Meissner, & Gaab, 2014; Snowling, 1998). Consequently, DD is increasingly characterised as a disruption of rapid auditory processing (RAP). Lehongre and colleagues (Lehongre et al., 2011) tested the auditory steady state response (ASSR) of individuals affected by DD and a group of controls. ASSR reflects an intrinsic and stable property of the brain that is relevant for auditory processing (Baltus & Herrmann, 2015). Lehongre and colleagues (2011) found that the magnitude of left lateralised ASSR to 30Hz amplitude-modulated stimuli was reduced compared to controls, and that the magnitude of the ASSR in planum temporale at 30Hz in both DD and control participants predicted performance on phonological tasks. They take this to be a reflection of an intrinsic property of the auditory system and conclude that an abnormal oscillatory rhythm in left auditory areas in the range 25-35Hz is responsible for DD. This demonstrates the relevance of this frequency range at this cortical area, for speech perception, consistent with our finding that RS activity in this frequency region at this anatomical locus predicts WIN performance. Further, deviance from normal ASSRs in the 25-35Hz range were observed by Lehongre and collaborators also in the left prefrontal cortex and in the right posterior superior temporal cortex, showing a high spatial similarity with regions that were correlated with WIN recognition abilities in our sample of healthy participants. Unfortunately, the boundaries of this frequency range exactly matches the centre of two different frequency bands defined in this study (21-29Hz and 30-40Hz, respectively), slightly complicating the comparison. Nevertheless, the ASSR is known to peak higher in the RH than LH (Lehongre et al., 2011; Ross, Herdman, & Pantev, 2005), and the effect we oserve may be related to this asymmetry. Indeed, it is possible that our results indicate that higher power at the peak ASSR frequency of the auditory system at rest is beneficial for speech perception. Under the assumption that the human auditory system has co-evolved with speech and is thus highly tuned to the acoustic properties of the latter, it may not be too much of a stretch to suggest that higher resting-state activity at the peak of the local neural tuning may provide a superior neural context for speech stimulus processing, underpinning improved speech in noise perception.

The present study is, perforce, subject to a number of limitations. One issue is the task that was used to evaluate speech in noise perception, namely the Words-In-Noise test. This presents participants only with a series of unconnected monosyllabic words. Use of such a set precluded investigation of another oscillatory component that has recently been tied to speech comprehension, namely the θ band (Ding, Melloni, Zhang, Tian, & Poeppel, 2015; Peelle & Davis, 2012). According to a number of reports and the predictions of the AST, θ underpins the syllabic parsing based on envelope cues, which are neither relevant nor present in the WIN. Further investigations of individual differences in performance on noisy connected speech materials would be necessary to evaluate whether RS θ may also contribute to the comprehension of acoustically challenging speech under slightly more ecologically valid conditions.

A further limitation relates to the fact that audiometry tests such as Pure Tone Audiometry (PTA) or tests of speech recognition in quiet were not part of the battery of behavioural assessments which we used to deconfound the WIN score. As a consequence, we cannot rule out that the effects we observed relate to the perceptual bases of speech recognition, or even hearing abilities per se. Nevertheless, most tests of speech-in-noise recognition tests, including the WIN test one used here, are based on a dual-component model of hearing (dis)ability: ‘signal amplification/attenuation’ (or audibility) refers to pure-tone sensitivity, whilst ‘signal filtering/distortion’ refers to the ability to understand speech in a noisy background (Plomp, 1986). These two functions are thought to be independent, or to be very weakly coupled at best. Indeed, although a few studies did report a relationship between pure-tone audiometry (PTA) and speech-in-noise performance (Lutman, 1991; Tschopp & Züst, 1994), the majority of the reports confirmed the dissociation between the two components (e.g., Blandy & Lutman, 2005; Dubno, Dirks, & Morgan, 1984; Duquesnoy, 1983; Jerger, 1992). One recent study reported that – when controlling for age – whilst PTA predicted speech recognition in quiet, these two measures were not associated with speech-in-noise recognition in neither normally-hearing nor hearing-impaired individuals (Vermiglio, Soli, Freed, & Fisher, 2012). Similarly and most importantly, the WIN test itself was shown to be independent from PTA as well as from speech recognition in quiet in elderly with mild-to-moderate hearing loss (Wilson et al., 2005), suggesting that threshold elevation in terms of signal-to-noise ratio, rather than signal attenuation, is what is chiefly assessed by this test. Thus, we believe that the associations between RS power and WIN recognition insofar described relate specifically to the perceptual foundations of resilience to noise-induced breakdown of speech intelligibility.

## 5. Conclusion

We have observed that analysis of spectrally-resolved RS-electrophysiology can provide valuable insights into the spatial and spectral properties of cerebral systems that are specific to WIN recognition. In this regard, the present results are consistent with the growing body of research indicating that the interactions between resting and evoked activity need to be better characterised in order to obtain a more complete and accurate account of cerebral functioning. In particular, beyond advancing our understanding of properties intrinsic to the brain that determine speech comprehension, a number of implications arise from this investigation. First, it suggests, together with a number of studies presented above, that altering resting oscillatory activity associated with RAP may constitute a viable approach for overcoming phonological processing deficits that arise from sub-optimal resting-state oscillatory activity patterns. This has already been shown to be the case within the realm of non-invasive electrical stimulation, but other interventions aimed at altering the balance of endogenous oscillatory activity associated with phonological processing can be conceived, such as EEG-informed neurofeedback, or closed-loop neurostimulation, targeting the spectral and spatial regions highlighted here. Furthermore, since the affordances of such interventions are inevitably tightly linked to pre-intervention activity levels, characterisation of RS activity in the normal population and in populations of patients affected by phonological impairments is a necessary step for the development of effective interventions. Thus, the scope of the analysis of neural correlates of RS power extends well beyond fundamental research and has important implications for applied research into how to support and enhance the auditory brain’s resilience in the face of noisy signals.

## Statement of Significance

We show that resting-state power, measured by MEG, in auditory cortices and left perisylvian areas is predictive of individual differences in speech in noise recognition performance. Intrinsic brain properties indexed by resting-state activity are behaviourally relevant, and provide further insight into the neural substrates of speech in noise processing capacity.

## Acknowledgements

Data were provided by the Human Connectome Project, WU-Minn Consortium (Principal Investigators: David Van Essen and Kamil Ugurbil; 1U54MH091657) funded by the 16 NIH Institutes and Centers that support the NIH Blueprint for Neuroscience Research; and by the McDonnell Center for Systems Neuroscience at Washington University.

The authors would like to thank three anonymous reviewers and Prof. P. Li for their constructive comments on an earlier version of the manuscript.

---

**Figure S1.**
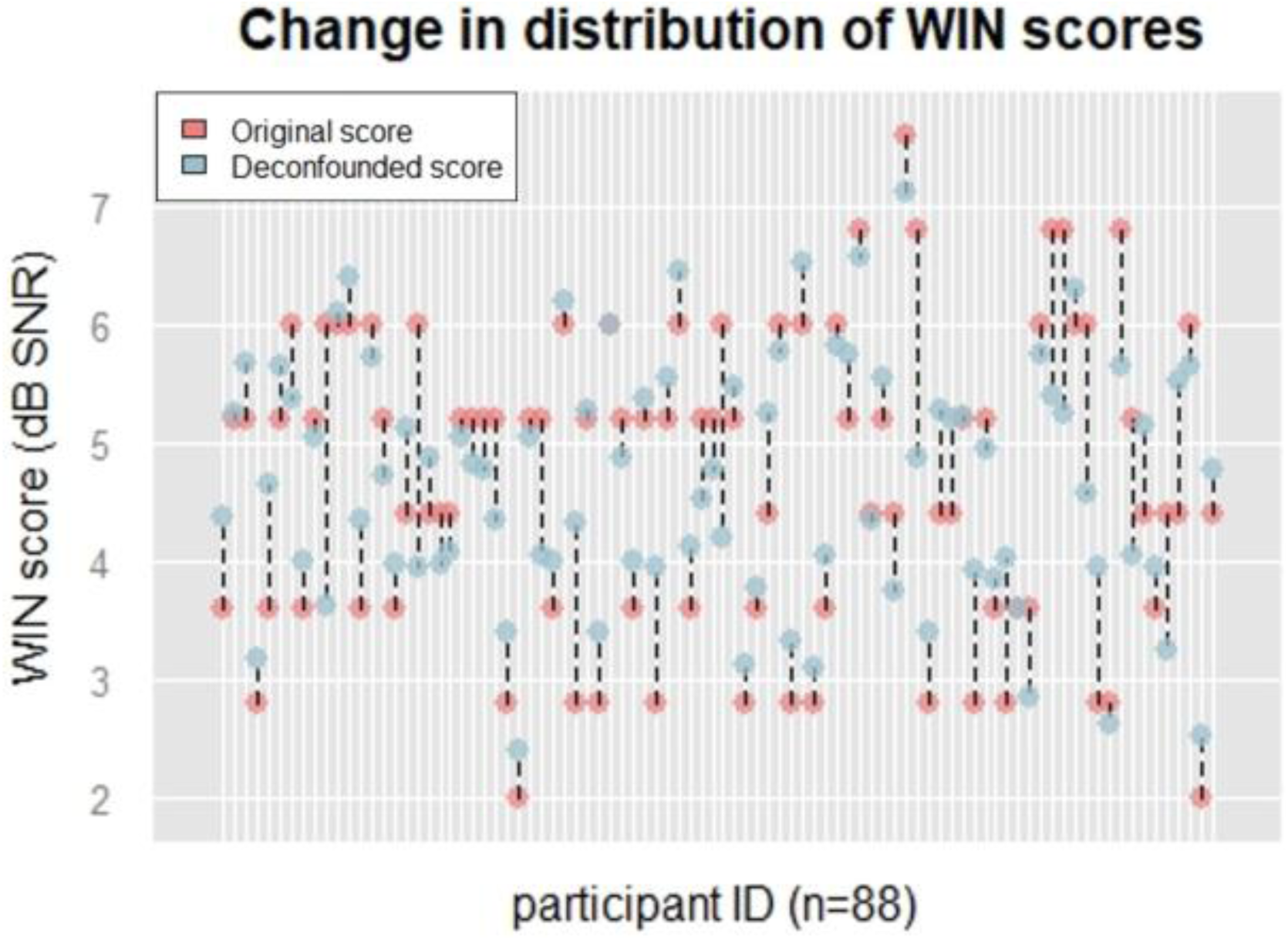
Change induced by deconfounding WIN on individual dB SNR scores. X-axis = participant ID; y-axis = WIN score in dB SNR; blue dots = original WIN scores; red dots = deconfounded WIN scores; dashed lines = change in WIN score associated with deconfounding procedure. Note: original WIN score was standardized to mean = 0 and SD = 1 before deconfounding; similarly, the deconfounded scores used to predict voxelwise power in the regression analysis were also standardized. Non-standardized scores are only us purposes.

**Figure S2.**
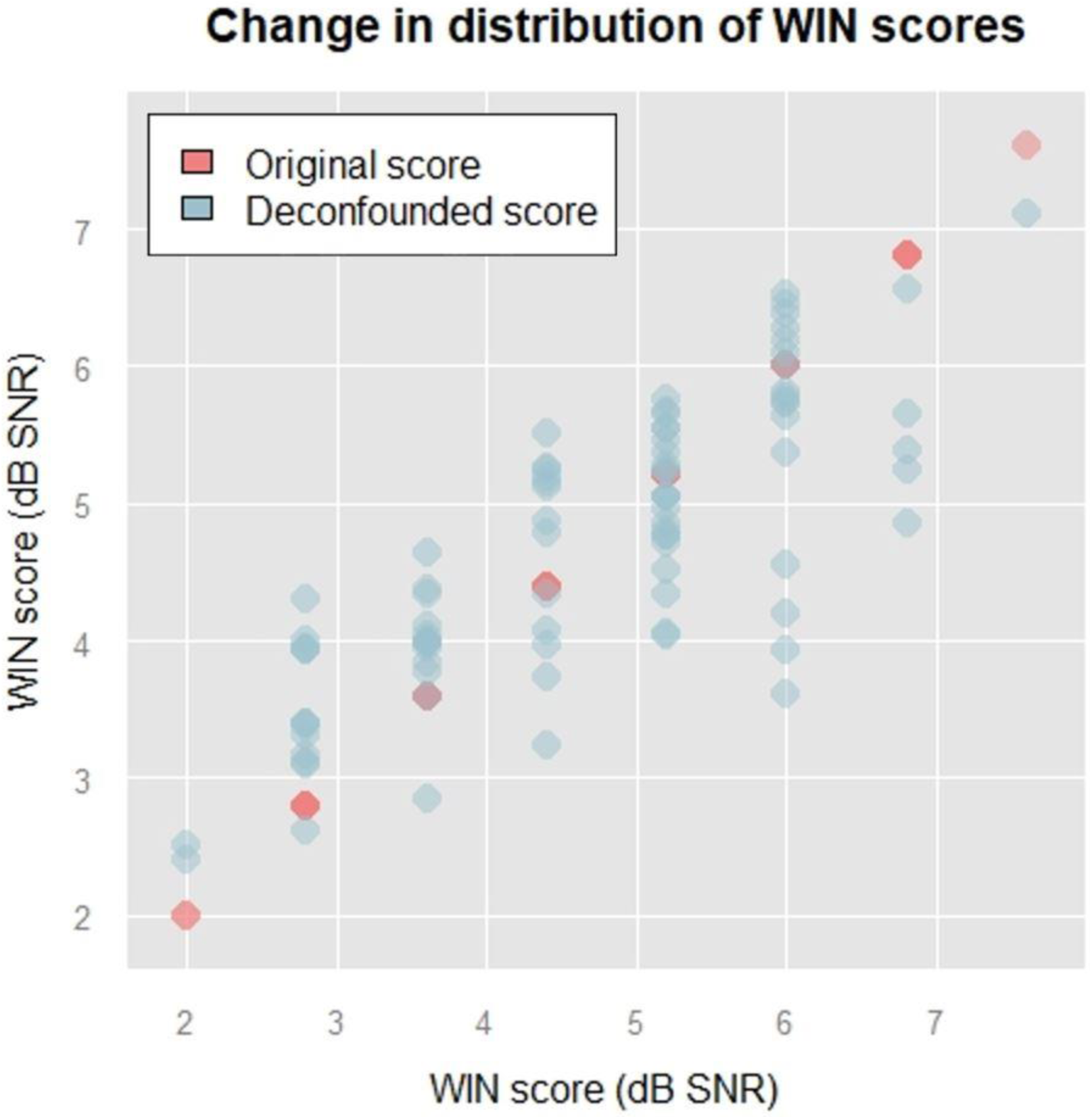
Effect of deconfounding WIN on the distribution of scores. Red dots represent the original scores (in dB SNR) of the corresponding subset of participants and blue dots above and below each original score represent each newly-deconfounded WIN score for each score-group. Performance of participants in our sample spanned eight dB SNR levels before deconfounding. The reduction in variance resulting from deconfounding is apparent in the way scores representing relatively better performance (2-4.4 dB SNR) increased on average (i.e., they represent relatively worse performance) and, vice versa, scores representing relatively worse performance (5.2-7.6 dB SNR) decreased on average (i.e., they represent relatively better performance).

## Appendix List and description of confounds.

*ID in* = identifier of the confounds before the selection procedure (see Table 1 for the selection criteria).

*Formal database name* = column headers of the “restricted” and “open access” csv files available here: http://humanconnectome.org/data.

*Full display name, Category, Assessment* = description of the confounds from the “data dictionary” csv file available here: https://wiki.humanconnectome.org/display/PublicData/HCP+Data+Dictionary+Public-+Updated+for+the+1200+Subject+Release?preview=/53444663/113377284/HCP_S1200_DataDictionary_April_20_2018.csv

*r, p(unc.), p(FDR)* = correlation coefficient, uncorrected p-value and FDR-corrected p-value of the correlation (Spearman’s rho) between the confound and the original (non-deconfounded) WIN SNR score. For three measures, these values are not available because the correlation could not have been computed as there was no variation of observations around the mean (same score for all f the participants).

*I* = indicates whether the measure has been included in the PCA (‘y’) or not (‘n’).

*ID out* = identifier of the selected confounds (those that entered the PCA).

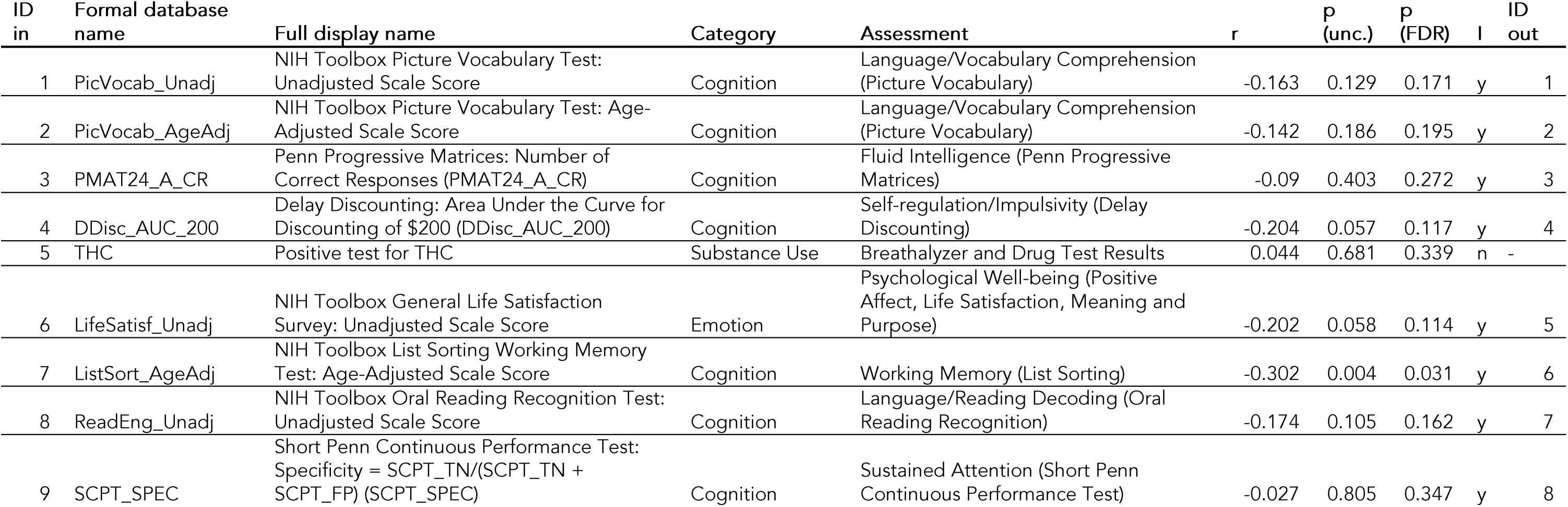
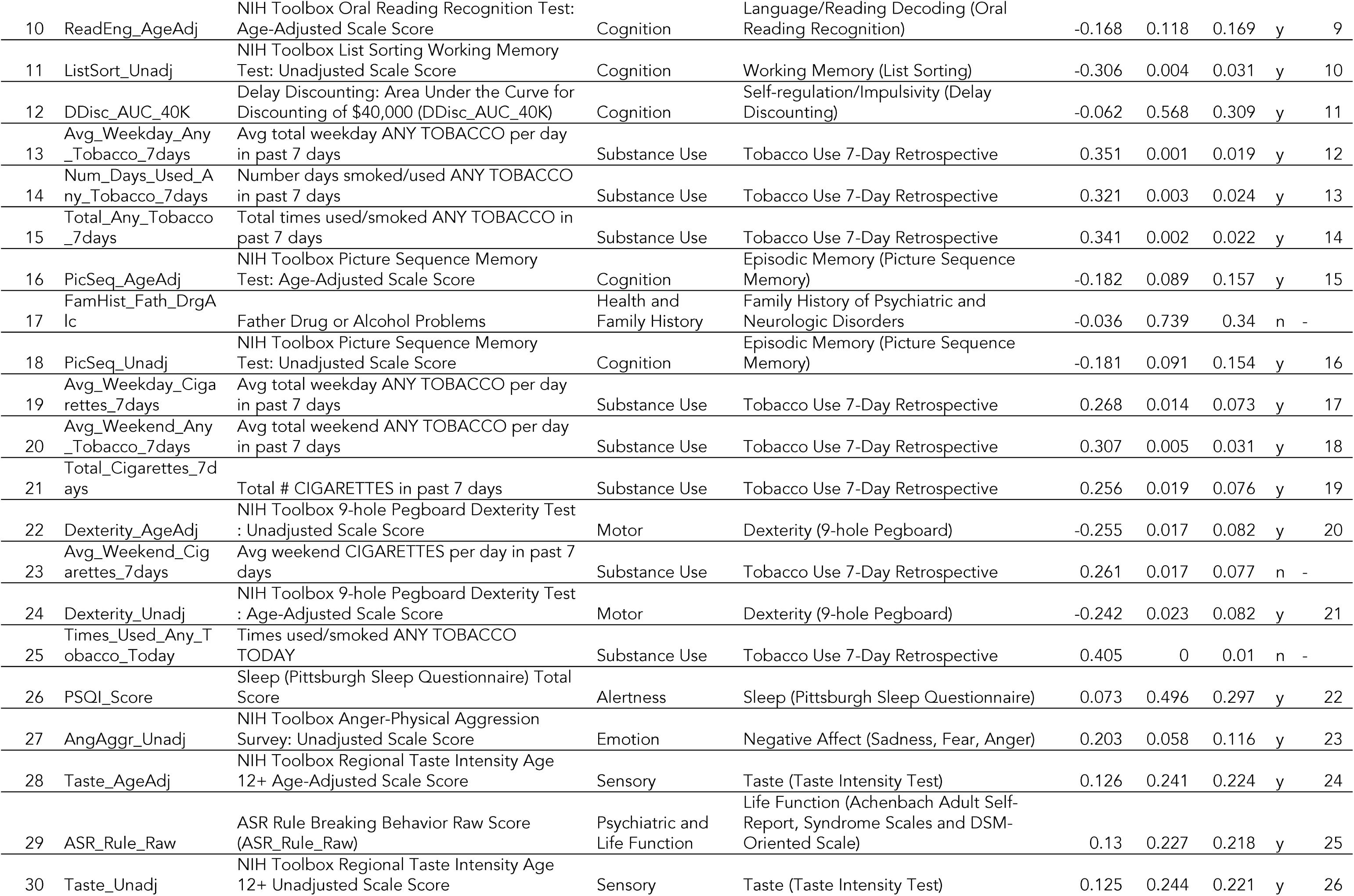
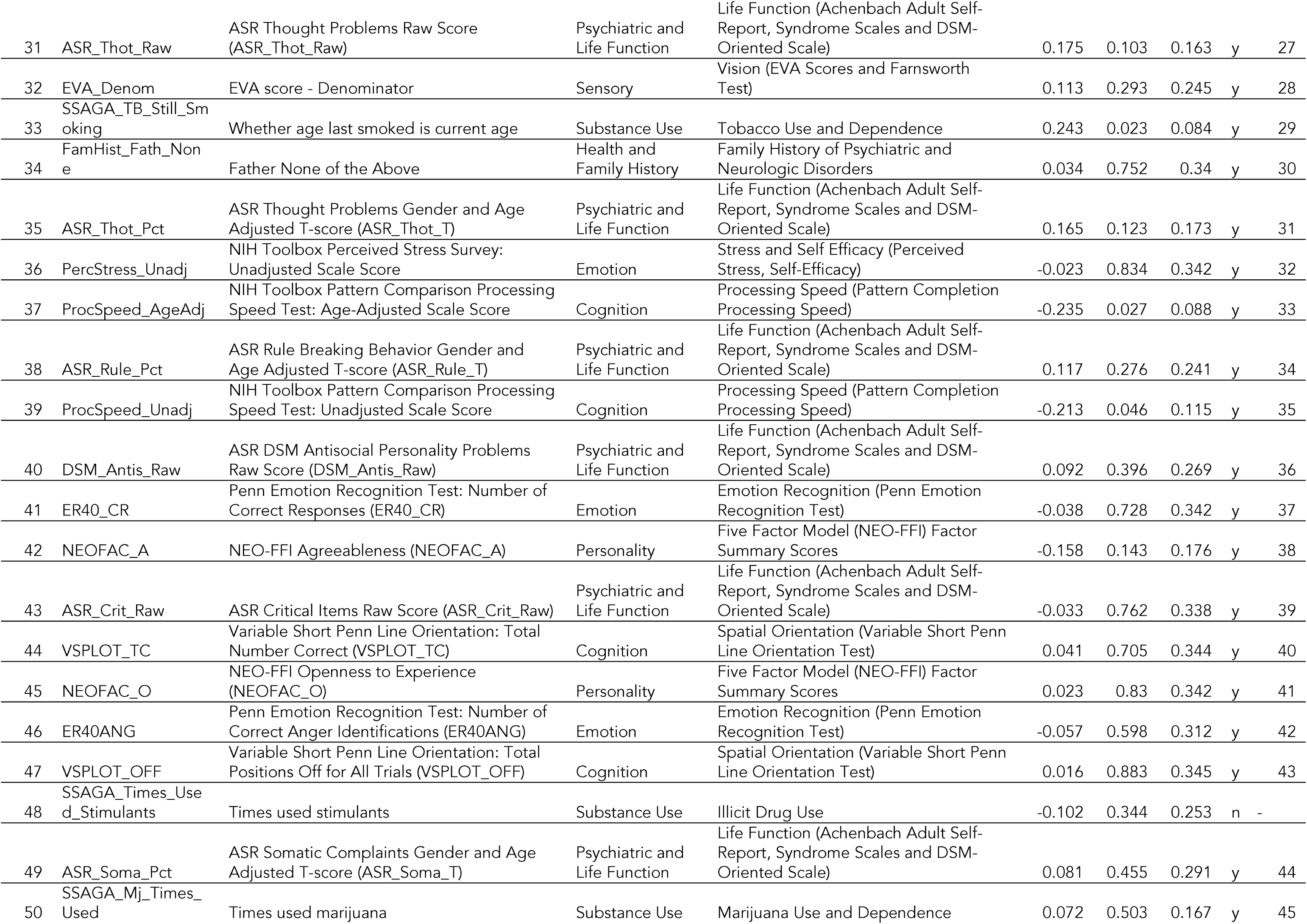
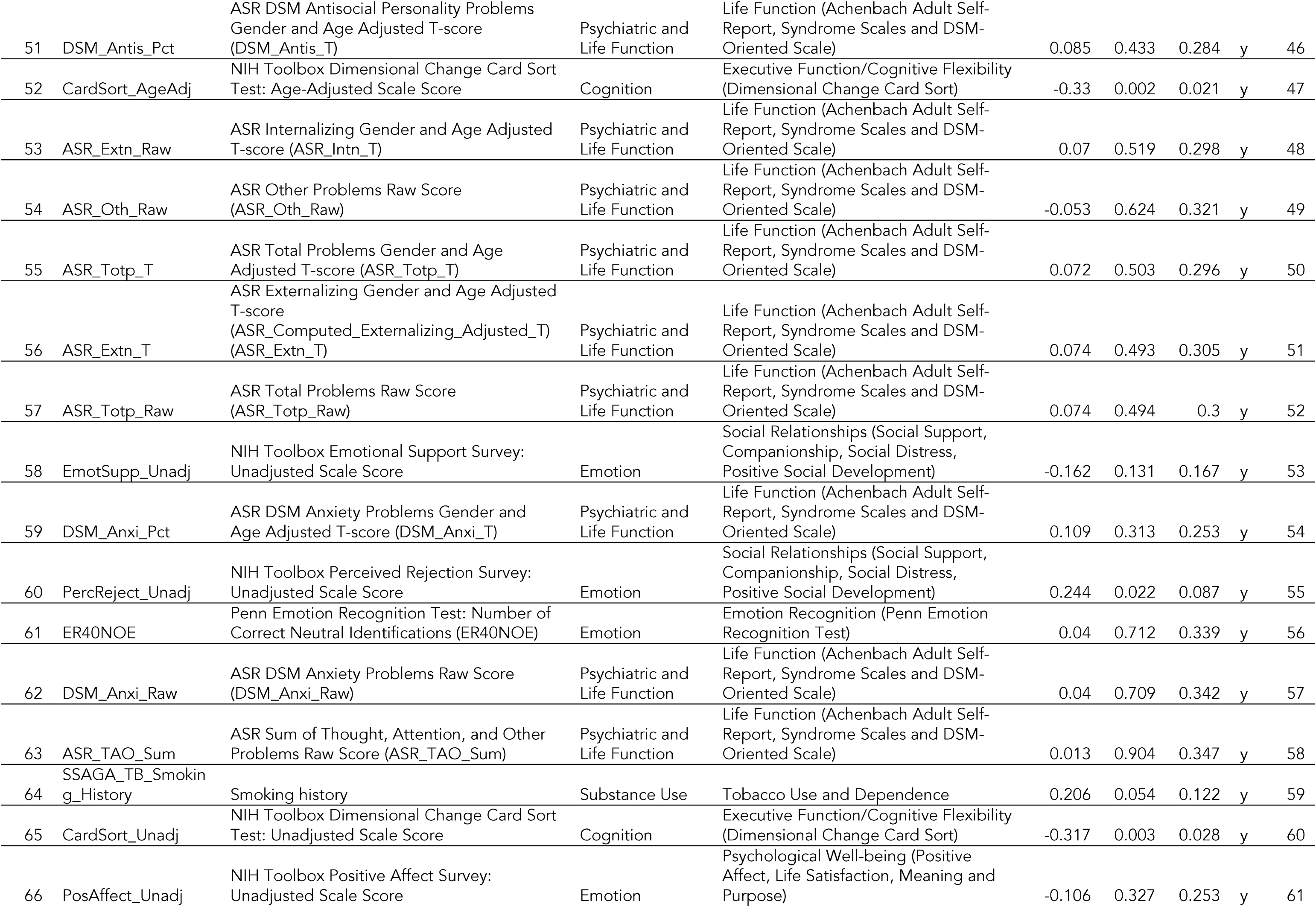
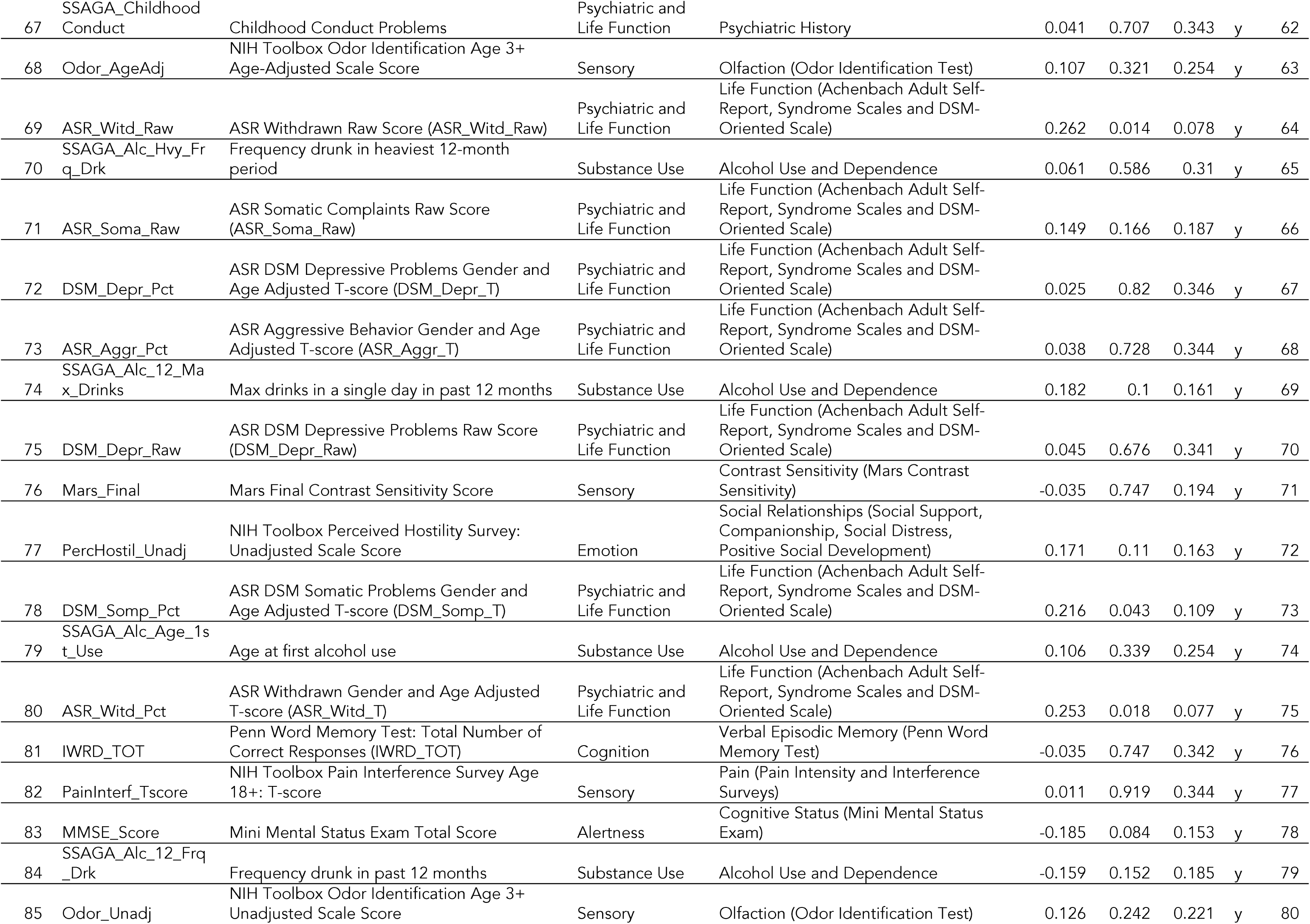
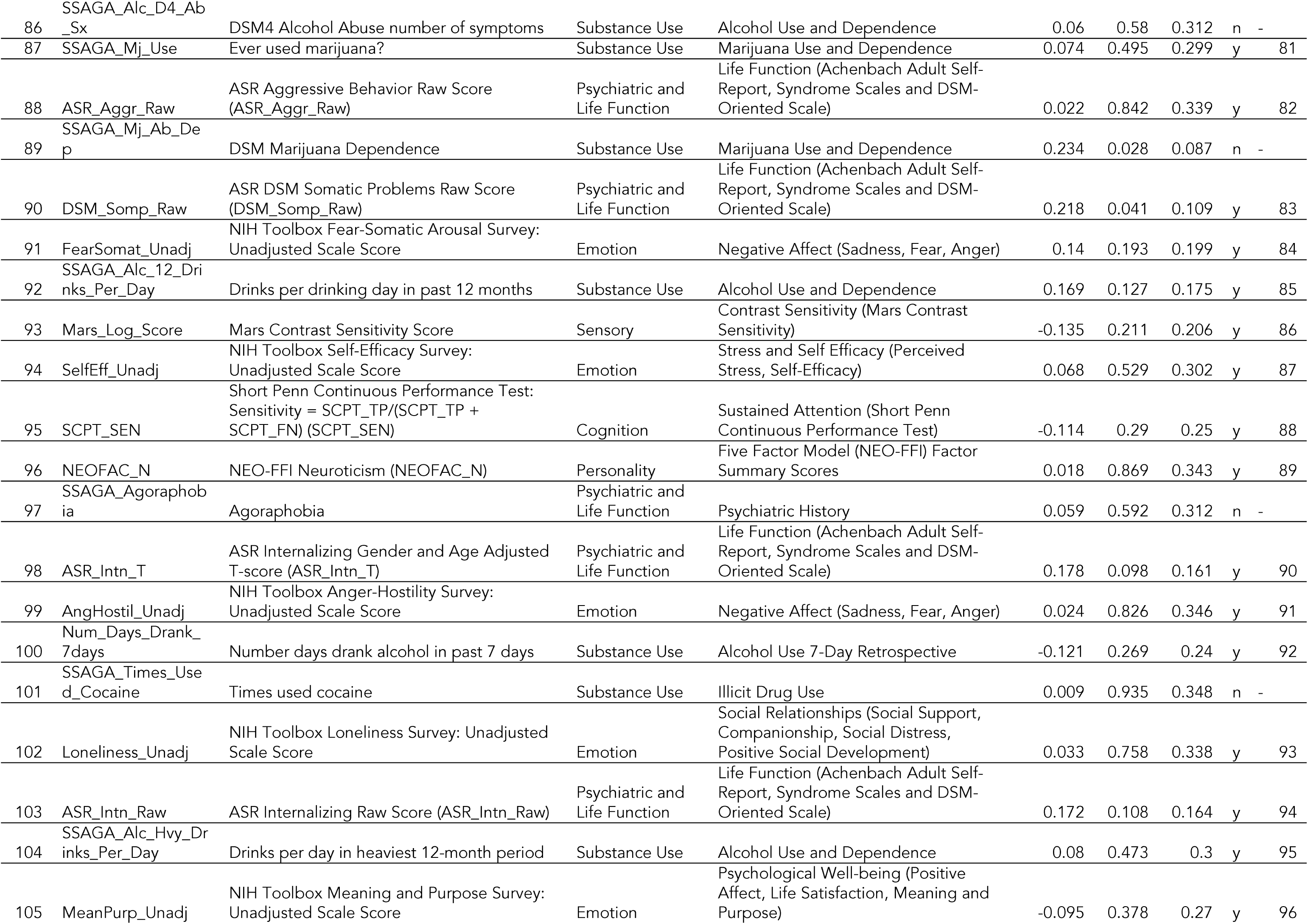
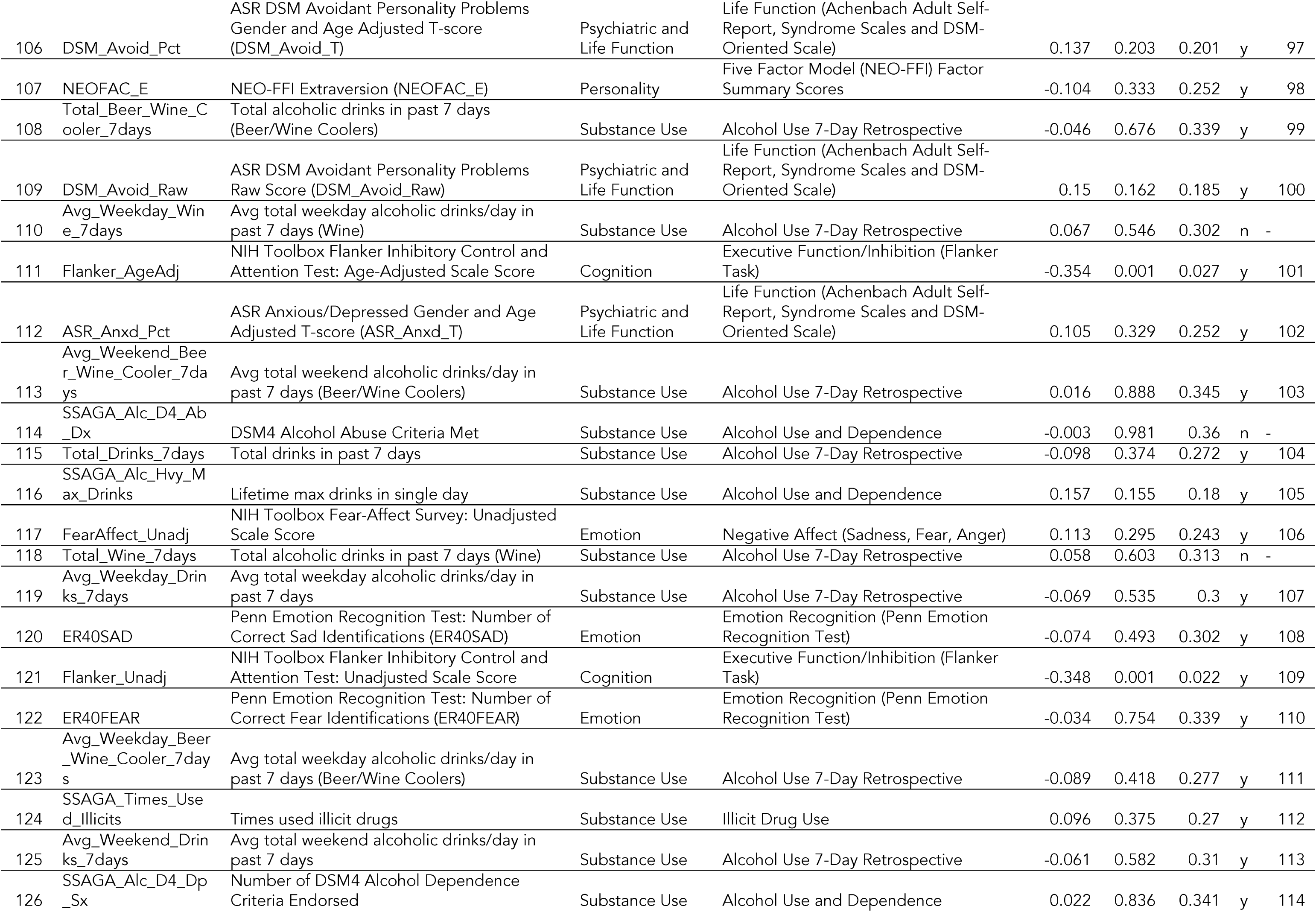
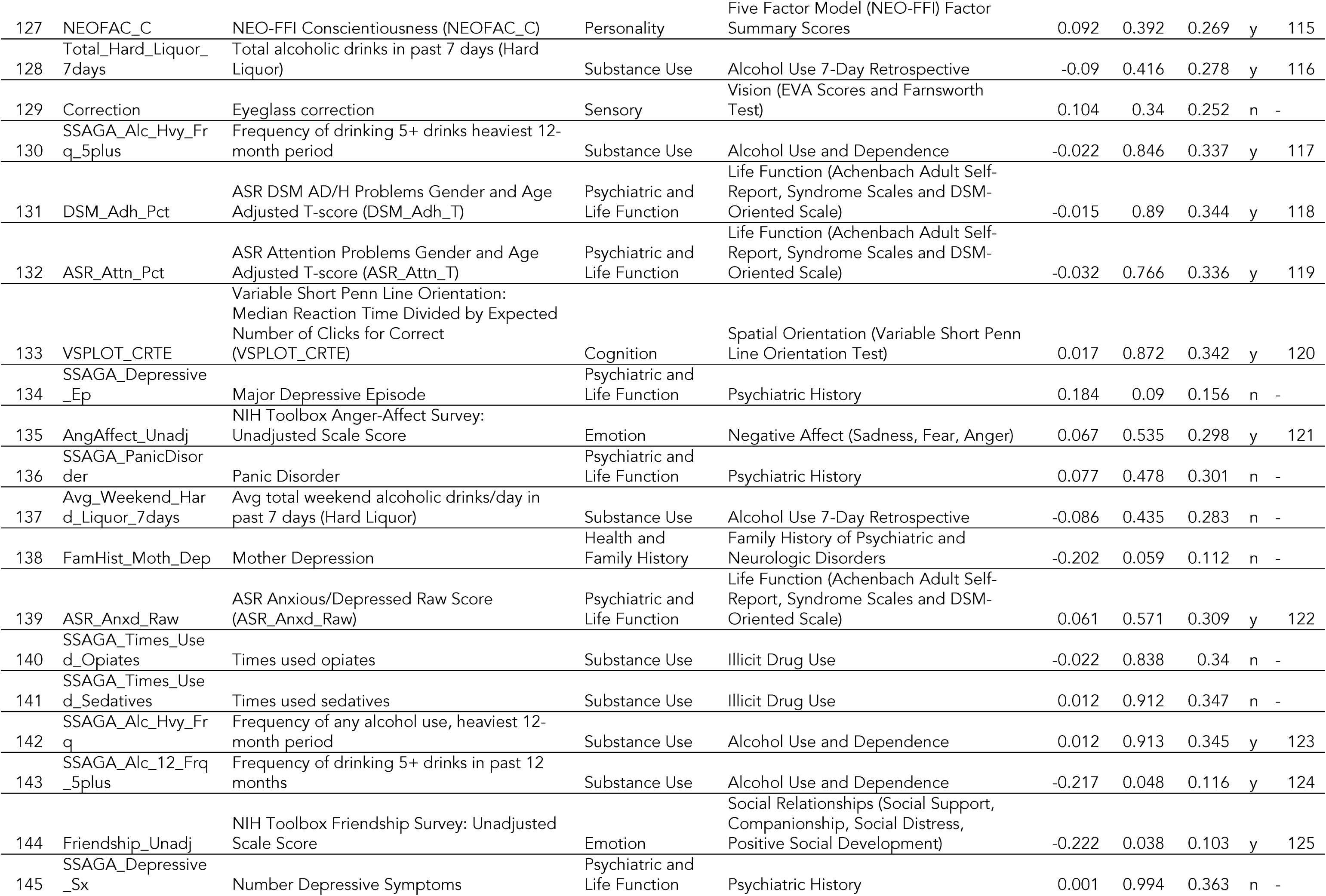
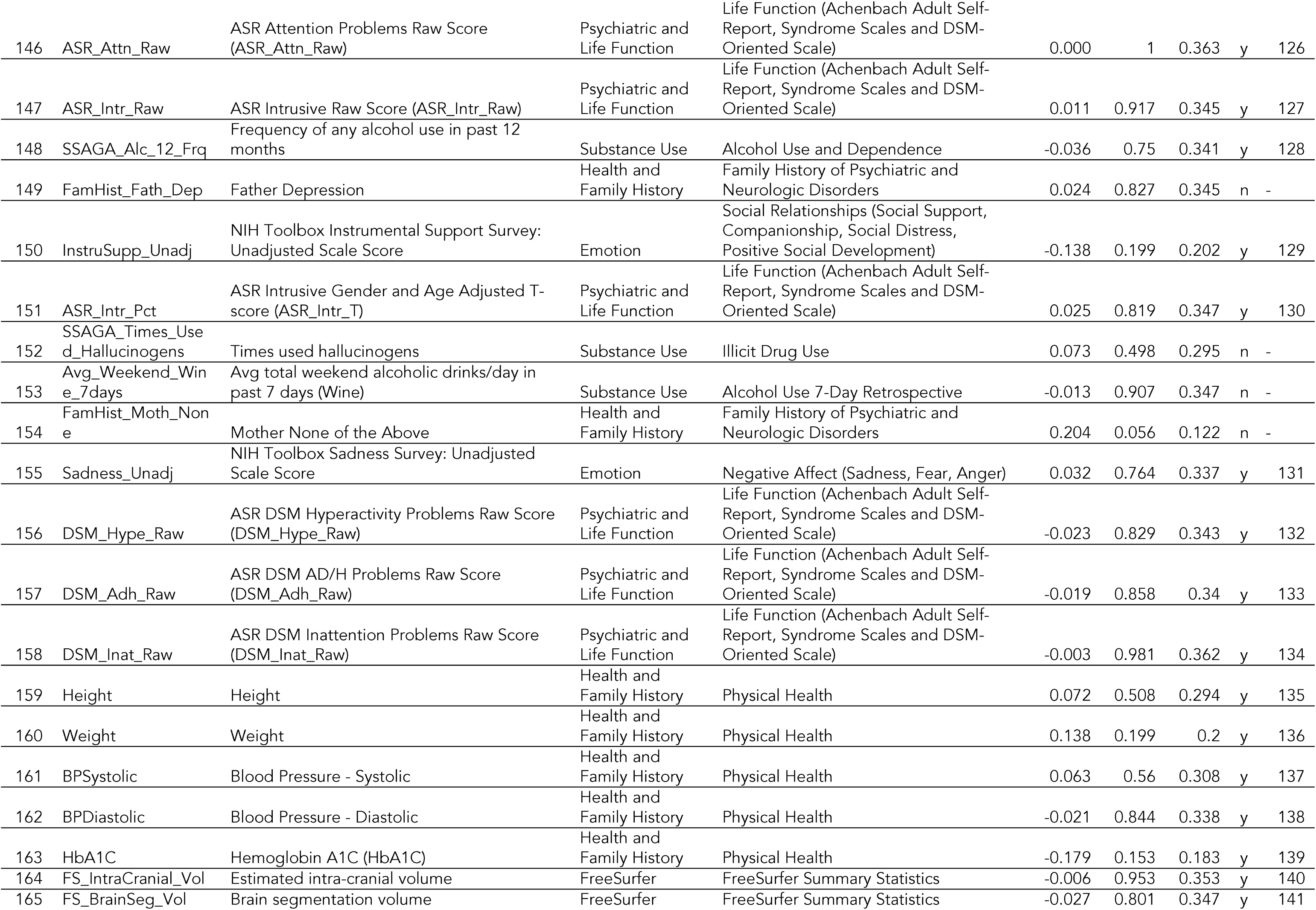
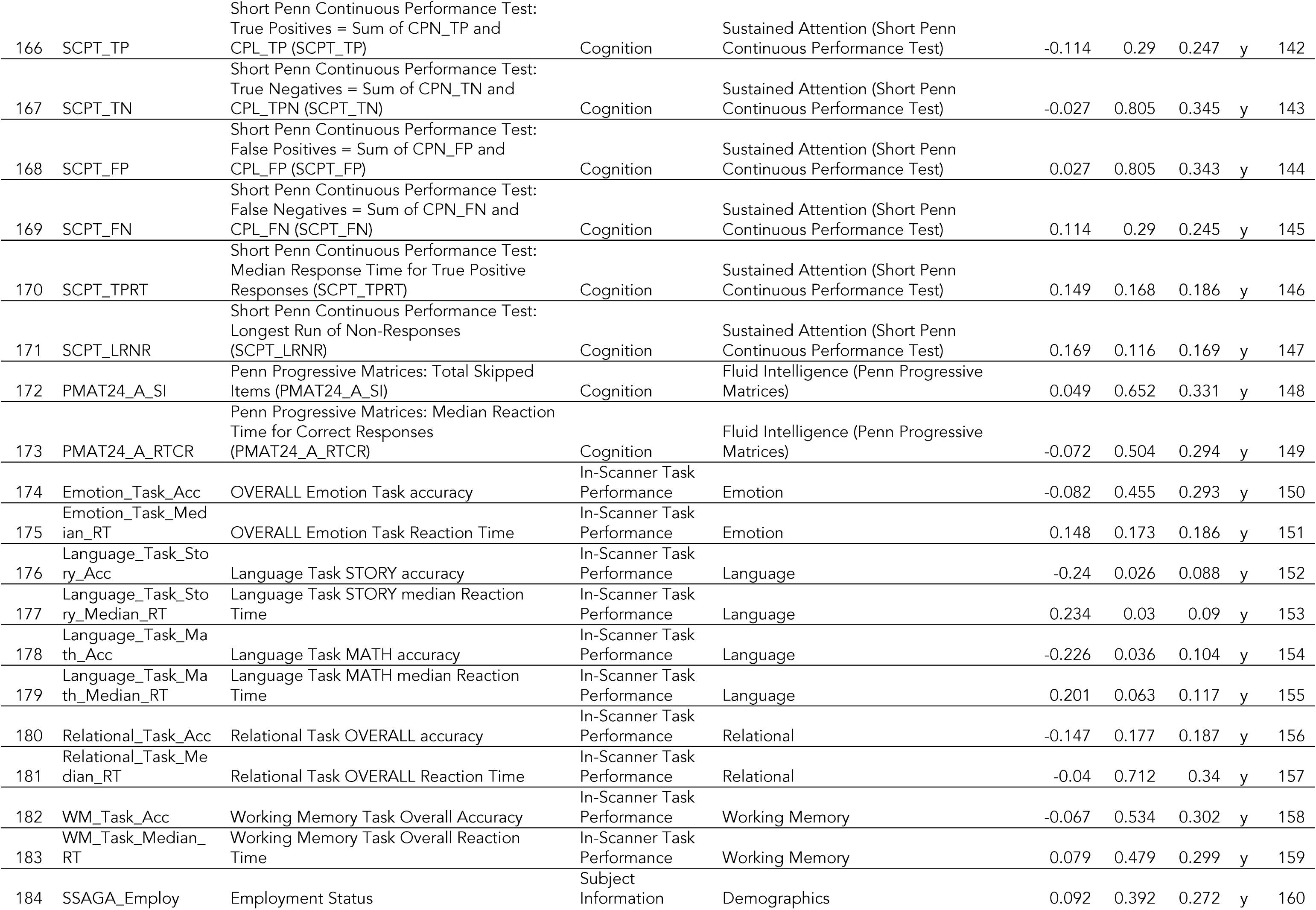
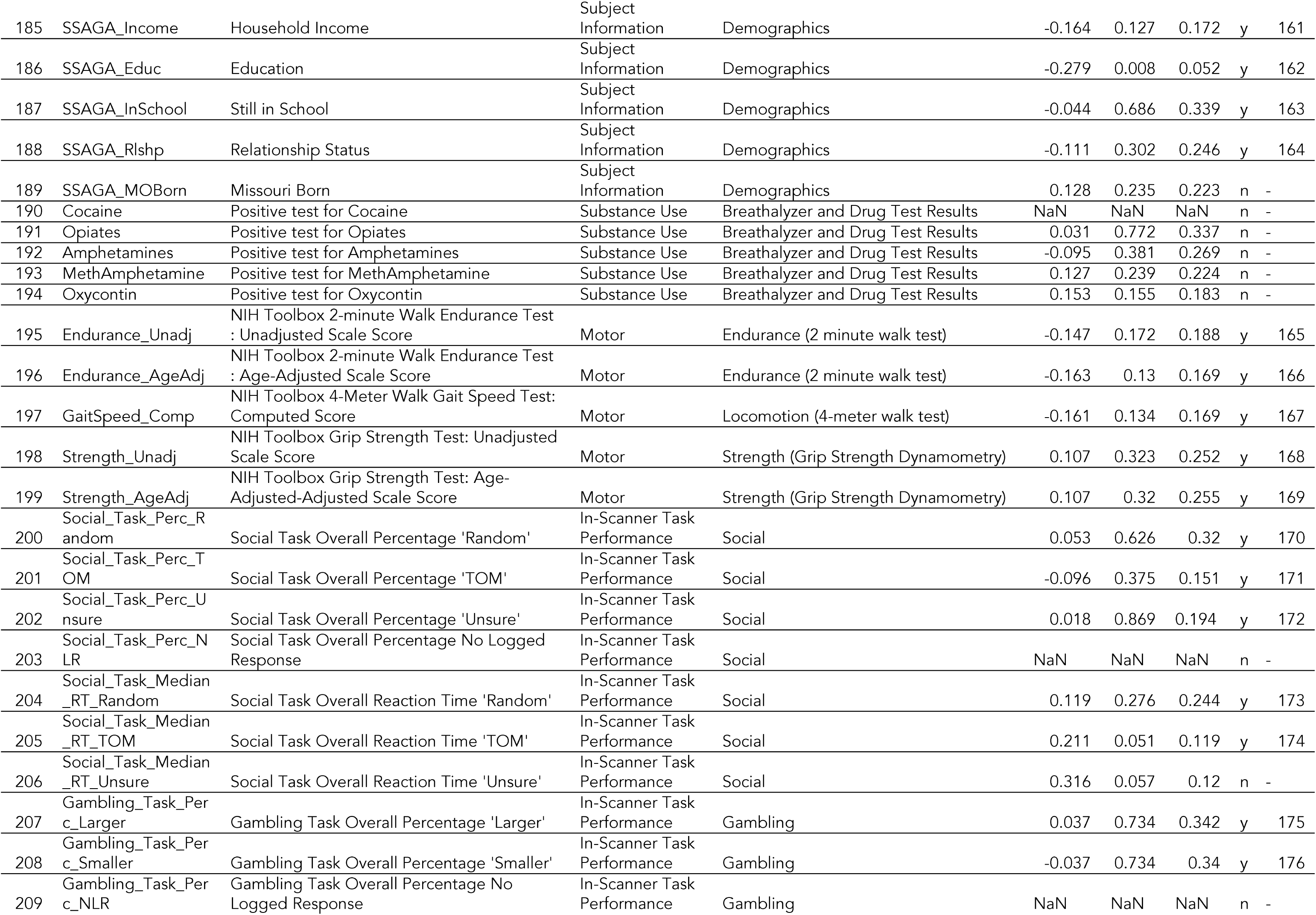
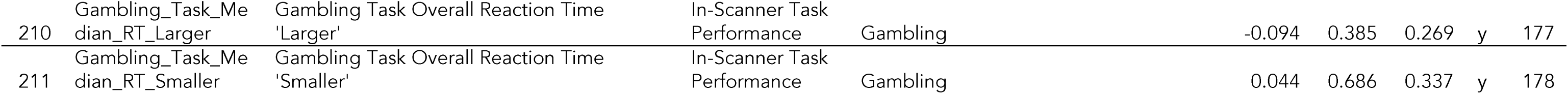

